# Cryogenic non-invasive 3D X-ray phase-contrast imaging of unfixed, intact mouse joints reveals shifting chondrocyte hypertrophy across the endochondral interface

**DOI:** 10.1101/2024.02.19.580961

**Authors:** L. A. E. Evans, D. Vezeleva, A.J. Bodey, P. D. Lee, G. Poologasundarampillai, A. A. Pitsillides

## Abstract

**Objectives:** i) develop and use a new cryogenically-enhanced phase contrast method to visualise hyaline articular cartilage (HAC); ii) to measure HAC, articular calcified cartilage (ACC) and total articular cartilage thicknesses in male STR/Ort (osteoarthritis, OA) and CBA (healthy) mouse tibial epiphyses, reflecting divergent OA predisposition, at three age timepoints chosen to reflect pre-OA, OA onset and late-progression; iii) to compare HAC, trans-zonal and ACC 3D chondrocyte anatomy in tibial epiphyses.

**Methods:** STR/Ort and CBA mouse knees (n=4 per age and strain group) were synchrotron-CT scanned at high-resolution while fresh frozen, without staining, fixation, dissection or dehydration of the joint capsule. Both cartilage thickness and cellular characteristics (chondrocyte n=420) were manually measured and statistically compared (SPSS).

**Results:** Cryo-enhanced phase contrast allowed cartilage to be seen in full thickness with cellular detail. HAC was thicker in STR/Ort than age-matched CBA mice in 16/24 knee joint compartments and timepoints (all p<0.04). In contrast, HAC was thicker only in the posterior lateral femur of CBA mice at 10weeks (p<0.001, Table 1). ACC and total cartilage were also thicker in STR/Orts. Trans-zonal chondrocytes were smaller than ACC and HAC chondrocytes (p-values<0.001, volumes 878, 1,567μm^3^ and 1,348μm^3^ respectively).

**Conclusions:** Cryogenically-enhanced phase-contrast imaging allowed cellular detail to be seen in 3D as never before in HAC in this (or any other) model. Our findings challenge current understanding by associating STR/Ort OA vulnerability with regions of thick, rather than thinning-with-age, cartilage. Our data affirm an association between excessively hypertrophic chondrocytes and OA is present in STR/Ort mice.

## Introduction

Osteoarthritis (OA) has substantially higher prevalence in adults >65 than <44 years^(1)^. No biomarker can reliably differentiate between young people who will age healthily and those in pre-OA stages who will develop clinical OA^(2, 3)^. Biomarker studies are therefore most valuable when longitudinal/cross-sectional, as predictors of onset or progression may be selectively expressed at certain points in the disease course^(4)^.

Animal models are often necessitated for such research, as these yield a predictable, compressed and repeatable timeframe of onset and progression, allowing confident comparison between groups^(5, 6)^. One such model for knee OA, accounting for 83% of human OA disability burden, is the inbred STR/Ort mouse in which males develop cartilage lesions and other OA characteristics at skeletal maturity^(6-8)^. This allows early (18-20 weeks) and late-stage OA (>32 weeks), and even pre-OA (<16 weeks) to be researched, without invasive imaging required to confirm pathology^(7)^.

Articular cartilage thickness is frequently measured to monitor joint health, directly from histology, confocal laser scanning microscopy or MRI, or indirectly, where it contributes to joint space width (JSW)^(9-17)^. Likewise, quantification of chondrocyte morphology and volume can be informative. When a chondrocyte’s phenotype becomes increasingly fibroblastic, alterations in metabolism and collagen synthesis may co-occur with shape changes^(18)^. Evaluations should accommodate the fact distinct knee compartments experience different locomotory loading patterns, and may therefore respond differently to stimuli and/or medications. This increases the value of reporting clear spatial descriptors where cartilage thickness is evaluated^(19)^.

Histology yields measurement of cartilage thickness and chondrocyte morphology in high-resolution, but is limited to 2D and conceivably by tissue processing artefacts^(20-22)^. Micro-computed X-ray tomography (μCT) offers a 3D alternative, but studies of soft tissues remain limited by low attenuation (giving low contrast); chondrocytes have been faintly observed in isolated bovine plugs, but only after phosphotungstic acid or ethanol submersion^(23)^. To address this, contrast-enhancing CT techniques emerged to increase soft tissue visibility by staining with chemicals including Lugol’s solution, Wells-Dawson polyoxometalate, osmium tetroxide or phosphotungstic acid^(24-27)^; propagation-based phase contrast with a partially-coherent beam^(28)^; and recently, cryogenic-enhanced phase-contrast μCT on cardiac tissue^(21)^.

Contrast-enhancing methods have been used in hyaline articular cartilage (HAC), a thin layer with minimal radiopacity^(23, 29-31)^. This study seeks to reveal HAC cellular detail at high-resolution in 3D, without staining, dehydration or dissection of the joint capsule, and no removal, decalcification or damage to the mineralised portion. This high-resolution 3D imaging of intact joints is sought via cryogenically-enhanced phase contrast synchrotron μCT (sCT) of OA (STR/Ort) and healthy (CBA, parental control strain) mouse knees at -20°C. Histological studies have found HAC in STR/Orts is thicker than in healthily-ageing CBAs^(13, 32)^. This study therefore also seeks to determine reproducibility of these findings by sCT.

Articular calcified cartilage (ACC) chondrocyte anatomy has been assessed in 3D using μCT^(29)^, however, it has not been evaluated in unfixed, non-dehydrated, non-stained HAC. STR/Ort mouse 3D chondrocyte characterisation is imperative because the OA may follow an inherent endochondral ossification defect^(32)^. One chondrocyte population in which any such defect may manifest, comprises cells potentially transitioning from hyaline-to-calcified articular cartilage zones across the tidemark (Evans et al.)^(33)^. This population, visible as ACC ‘pits’ when HAC is removed with proteases^(34, 35)^ – referred to hereafter as ‘trans-zonal’ – is, however, entirely uncharacterised in 3D in mouse joints. Our aims were therefore to: i) develop and use a new cryogenically-enhanced phase contrast method to visualise HAC; ii) measure HAC, ACC and total articular cartilage thicknesses in STR/Ort and CBA tibial epiphyses, reflecting divergent OA predisposition, at three ages chosen to reflect pre-OA, OA-onset and late-progression; and iii) compare HAC, trans-zonal and ACC 3D chondrocyte anatomy.

## Methods

### Animals

Procedures were performed in accordance with the Animals (Scientific Procedures) Act 1986. Study design was approved by Clinical Research Ethical Review Board at the Royal Veterinary College (RVC), and the United Kingdom Government Home Office under special project licence. Male STR/Ort or CBA mice were used.

STR/Ort were bred in the RVC Biological Services Unit via sibling mating and housed in single-sex cages, avoiding isolated housing. CBA (∼50-day-old) were purchased (Charles River) and aged in-house. Standard open cages were ventilated with 15-20 air changes/hour. Maximum number/cage is 5 (if mice weigh ≥20g) to 8 (if <20g).

Bedding comprised wood chip, nesting materials and paper, enriched with plastic and cardboard tunnels. 21°C (± 2°C) temperature was maintained with 12 hour light/dark cycles. Animals accessed water and standard RM1 maintenance diet (Special Diet Services, South Witham, UK) ad libitum. Mice were killed at 9-11, 18-20, or 36+ weeks of age (referred to hereon as 10wk, 20wk and 36+wks) by cervical dislocation or rising CO_2_; ages correspond with pre-OA, OA-onset and late-OA in STR/Ort mice. Mean body weight at death was 34g with no significant strain difference.

### Sample preparation

Knees were scanned with joint capsule intact (see Supplementary Tables 1-2 for cartilage thickness and chondrocyte evaluation sample sizes). Hindlimbs were dissected, skinned, and musculature removed. Rectus femoris was left to maintain patellofemoral integrity. Proximal tibia and distal femur were cut by scalpel to keep articular surfaces intact, with bones extending ∼1.5mm proximally and distally. Knees were wrapped in phosphate-buffered saline-soaked gauze, stored in plastic bijous, and frozen at -80°C. Bijous were transported to Diamond Light Source (DLS) on dry ice, and transferred to -20°C freezer storage on arrival.

Fresh frozen samples were parafilm-wrapped and placed into 5mm-diameter Kapton tubing, affixed to holders. To control temperature and reduce motion, a cold stage (Guo et al. (2017); developed specifically for DLS I13-2 use), was mounted around the sample holder, enabling temperature control remotely (Supplementary Fig. 1)^(36)^ to -20±3°C.

### Scanning and reconstruction

Samples were scanned at I13-2 (DLS) using a filtered pink beam of 27 keV weighted-mean photon energy (5 mm undulator gap; Pt mirror stripe; filters: 1.3 mm graphite, 3.2 mm Al) and a pco.edge 5.5 CL camera (PCO AG, Germany) mounted on a scintillator-coupled visible-light microscope of variable magnification. Up to 4001 projection images were acquired per 180° scan with 0.05 s exposure time, 1.625 µm voxel size and 4.2 × 3.5 mm^2^ (2560 × 2160 pixels) field of view. The first (0°) and last (180°) images were compared to check for scanning errors^(37)^. Sample-detector distance was adjusted to alter extent of phase contrast. Scans were reconstructed via filtered back projection in the modular pipeline Savu^(37, 38)^, incorporating flat- and dark-field correction, optical distortion correction^(39, 40)^, ring artefact suppression^(41)^ and automatic rotation centre calculation^(42)^.

### Exclusion

Although sample size originally included 6-8 limbs/group, final size was reduced (Supplementary Tables 1-2) by factors including: radiation damage, motion or ring artefacts, failure to capture full joint in FoV, and time-limited access. Groups were equalised (Supplementary Table 1) to minimise bias.

### Cartilage thickness

Image stacks were orientation-matched, resliced into coronal plane (Dataviewer, Bruker, Belgium) and opened in increments (30 slices, 48.75μm) in ImageJ^(43)^.

Cartilage thickness was measured in all condyles (medial tibia/femur, lateral tibia/femur) by drawing and recording the length of a vertical line at the condylar centre, identifiable by drawing a line horizontally across the articular surface (as centre of all lines drawn in ImageJ are marked, Fig. 1B). The central line of each condyle was initially drawn from the superficial surface to the mineralising front, to measure HAC thickness (Fig. 1C). The deep end of the same vertical line was then extended to the cement line (bone/calcified cartilage junction) (Fig. 1D), and re-measured, providing a second location-matched total cartilage thickness. On rare occasions HAC lesions were present (Fig. 1E), HAC and total cartilage thickness measurements were instead recorded at the nearest in-plane location where HAC was intact (Fig. 1F-G) and repeated until HAC and total cartilage thicknesses had been measured on all opened slices from posterior to anterior, for all condyles per knee.

**FIGURE 1:**
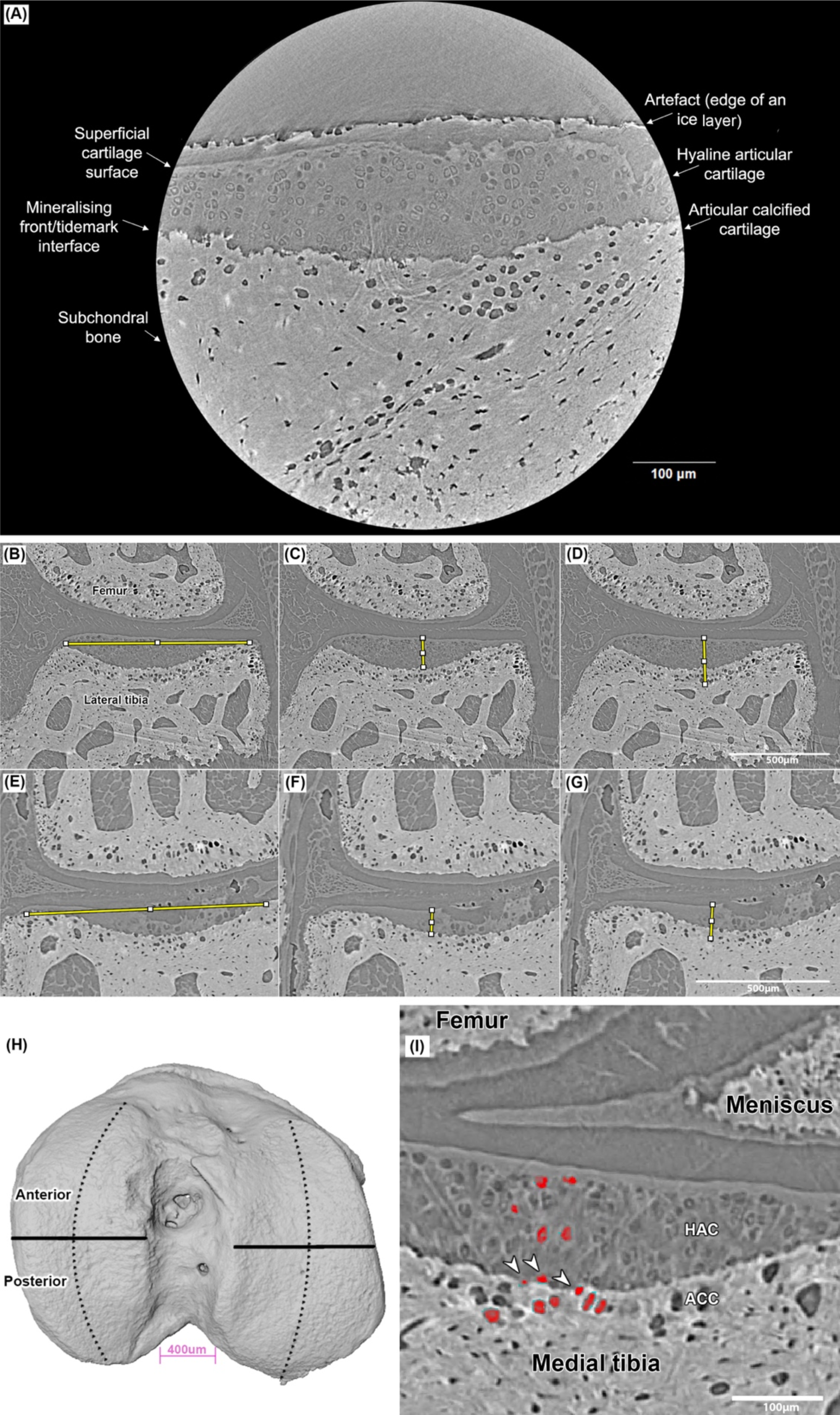
Phase contrast imaging of intact mouse knee joints using synchrotron computed tomography. (A) 40 week old male STR/Ort medial tibia (0.325μm pixels). Note that HAC and its resident chondrocytes are visible throughout, including within the superficial layer. (B-G) Method of measuring cartilage thicknesses on intact and lesioned tibial plateaus viewed in the coronal plane, using (B-D) a 10 week old STR/Ort mouse knee, in which (B) the centre of the condyle is identified by drawing a line horizontally across it (yellow, the centre is depicted as a small white square; (C) HAC depth is measured and (D) total articular cartilage depth is measured. (E-F) 36+ week old STR/Ort mouse, in which (E) the centre of the condyle is identified but HAC is visibly eroded in that location, therefore (F) HAC depth is measured in the nearest lesion-free region, as is (G) total cartilage depth. (H) Attributing locality to articular cartilage thickness measurements; 10 week old CBA tibial epiphysis rendered in 3D using Avizo. Dotted black lines represent the central axis of each condyle, from which serial cartilage thickness measurements were taken (medial condyle on the left, lateral condyle on the right). Solid black lines represent the central divide of each condyle, on either side of which thickness measurements were deemed ‘anterior’ or ‘posterior.’ (I) Example of a medial tibial plateau from which 15 chondrocytes were segmented (10 week old STR/Ort). Hand-segmented chondrocytes are highlighted in red; the three (out of five total) visible trans-zonal chondrocytes are identified by white arrowheads. Note that femoral HAC (visible at the top-left) is too thin to reveal cellular detail, and neither does the most superficial tibial HAC layer.

Measurements were spreadsheet-annotated with mouse strain, age, unique sample number and anterior or posterior joint location. Each HAC thickness measurement was subtracted from the total thickness measurement at the corresponding joint slice to yield ACC thickness. HAC and ACC thicknesses were also converted into relative values (percent total). Sometimes ACC comprised 100% of cartilage; these data were deleted to ensure measurements were recorded only from the articular condylar region (calcified cartilage was observed to extend anteriorly over the epiphysis).

### Chondrocyte manual-segmentation

Hand-segmentation of individual HAC, ACC and trans-zonal chondrocytes was completed while blinded (Supplementary Methods). Key criteria were that chondrocytes must have: i) clearly defined edges; ii) no contact with any nearby cell, or lacunae merging into those of any others; iii) be entirely within a chosen zone (applies only to HAC and ACC chondrocytes; eg. a HAC chondrocyte must not contact the tidemark, but be entirely HAC-surrounded; Fig. 1I). Medial and lateral tibial condyles were evaluated.

Femoral HAC was generally too thin to resolve HAC and trans-zonal chondrocytes (Fig. 1I). HAC and trans-zonal chondrocytes also tended not to be visible in CBA mice, due perhaps to cartilage thinness (Table 1), reducing sample size. The final sample of hand-segmented chondrocytes numbered 420; knee joints from which 30 chondrocytes each were segmented are in Supplementary Table 2.

**TABLE 1:** Condylar compartments in which HAC thickness was measured, and the presence or absence of a significant difference between STR/Ort and CBA mouse strains. * indicates a significant difference between strains (p<0.05). Note that the posterior lateral femur at 10 weeks of age is the only region in which STR/Ort mice have significantly thinner HAC than CBA mice.

| Age (weeks) | Condyle | Anterior/Posterior | P value for strain difference | CBA HAC thickness (µm) | STR/Ort HAC thickness (µm) |
| --- | --- | --- | --- | --- | --- |
| 10 | Lateral femur | Anterior | 0.672 | 20.8 | 19.5 |
|  |  | Posterior | **< 0.001\*** | **26.0** | **19.0** |
|  | Lateral tibia | Anterior | 0.223 | 14.6 | 13.1 |
|  |  | Posterior | **< 0.001\*** | **50.4** | **80.8** |
|  | Medial femur | Anterior | **< 0.001\*** | **17.1** | **26.1** |
|  |  | Posterior | **< 0.001\*** | **20.1** | **27.6** |
|  | Medial tibia | Anterior | 0.113 | 19.0 | 25.8 |
|  |  | Posterior | **< 0.001\*** | **43.0** | **87.1** |
| 20 | Lateral femur | Anterior | **0.019\*** | **17.9** | **23.1** |
|  |  | Posterior | 0.247 | 26.3 | 21.6 |
|  | Lateral tibia | Anterior | **0.008\*** | **13.0** | **14.6** |
|  |  | Posterior | **< 0.001\*** | **43.9** | **65.6** |
|  | Medial femur | Anterior | **< 0.001\*** | **17.5** | **32.3** |
|  |  | Posterior | **< 0.001\*** | **19.9** | **39.0** |
|  | Medial tibia | Anterior | **< 0.001\*** | **16.1** | **26.1** |
|  |  | Posterior | **0.038\*** | **47.2** | **71.5** |
| 36+ | Lateral femur | Anterior | 0.396 | 19.6 | 21.3 |
|  |  | Posterior | 0.549 | 27.6 | 24.4 |
|  | Lateral tibia | Anterior | 0.193 | 13.4 | 10.9 |
|  |  | Posterior | **< 0.001\*** | **40.9** | **58.1** |
|  | Medial femur | Anterior | **< 0.001\*** | **14.5** | **31.4** |
|  |  | Posterior | **< 0.001\*** | **16.6** | **35.2** |
|  | Medial tibia | Anterior | **0.004\*** | **15.4** | **21.2** |
|  |  | Posterior | **< 0.001\*** | **39.1** | **76.6** |

### Statistics

Statistical analyses were performed in SPSS. Cartilage thickness data were neither normally distributed (tested by Kolgorov-Smirnov and Levene’s tests), nor normalisable by log, square-root and/or inverse transformation. Non-parametric Kruskal-Wallis test was therefore applied with 48 groups, taking into account strain (STR/Ort, CBA), age (10, 20, 36+wks), condyle (lateral femur/tibia, medial femur/tibia), and location (anterior or posterior). This yielded abundant data, much of which was non-significant or beyond the scope of our hypothesis. Thus only strain differences per subgroup (including at different ages) are reported.

Residuals of volume and surface area data derived from individual 3D analysis (i3D) were normalised through log10, while sphericity residuals were normalised through log10 followed by inverse transformation (Kolgorov-Smirnov, all p-values≥0.2). Linear mixed model analysis was applied with four fixed factors (strain; age; lacunar depth; condyle) and one random factor (individual ID, to take into account repeated measurements/mouse). Post-hoc two-way interaction effects were calculated between all fixed factors. Since sample sizes for different mouse groups varied (Supplementary Table 2), Bonferroni correction was applied throughout.

Residuals of chondrocyte orientation (theta) derived from i3D of hand-segmentation were neither normal, nor normalisable through data transformation (Kolgorov-Smirnov, all p-values<0.001). Non-parametric Kruskal-Wallis test was applied with 36 groups, taking into account strain, age, tibial plateau and cartilage depth (HAC, trans-zonal, ACC). For brevity and clarity, significant differences are reported only where any single variable differs between groups, because this is informative of which fixed factor is relevant to observed lacunar orientation differences. A post-hoc Bonferroni correction was applied to all p-values.

## Results

### Soft tissue visualisation

HAC and other soft tissues were visible in high-resolution with 3D cellular-level detail, without fixation, staining, dissection, invasive processing or dehydration, in fresh frozen joints imaged at -20°C. Sample-detector distance adjustments beyond 60mm did not visibly improve phase contrast (Fig. 2A-B). In contrast, tissue fixation obliterated the capacity to resolve cellular-level detail, even with other imaging settings replicated (Fig. 2C-D). Fixation also induced HAC lesion swelling, reducing cartilage fibrillation visibility (Fig. 2C-D).

**FIGURE 2:**
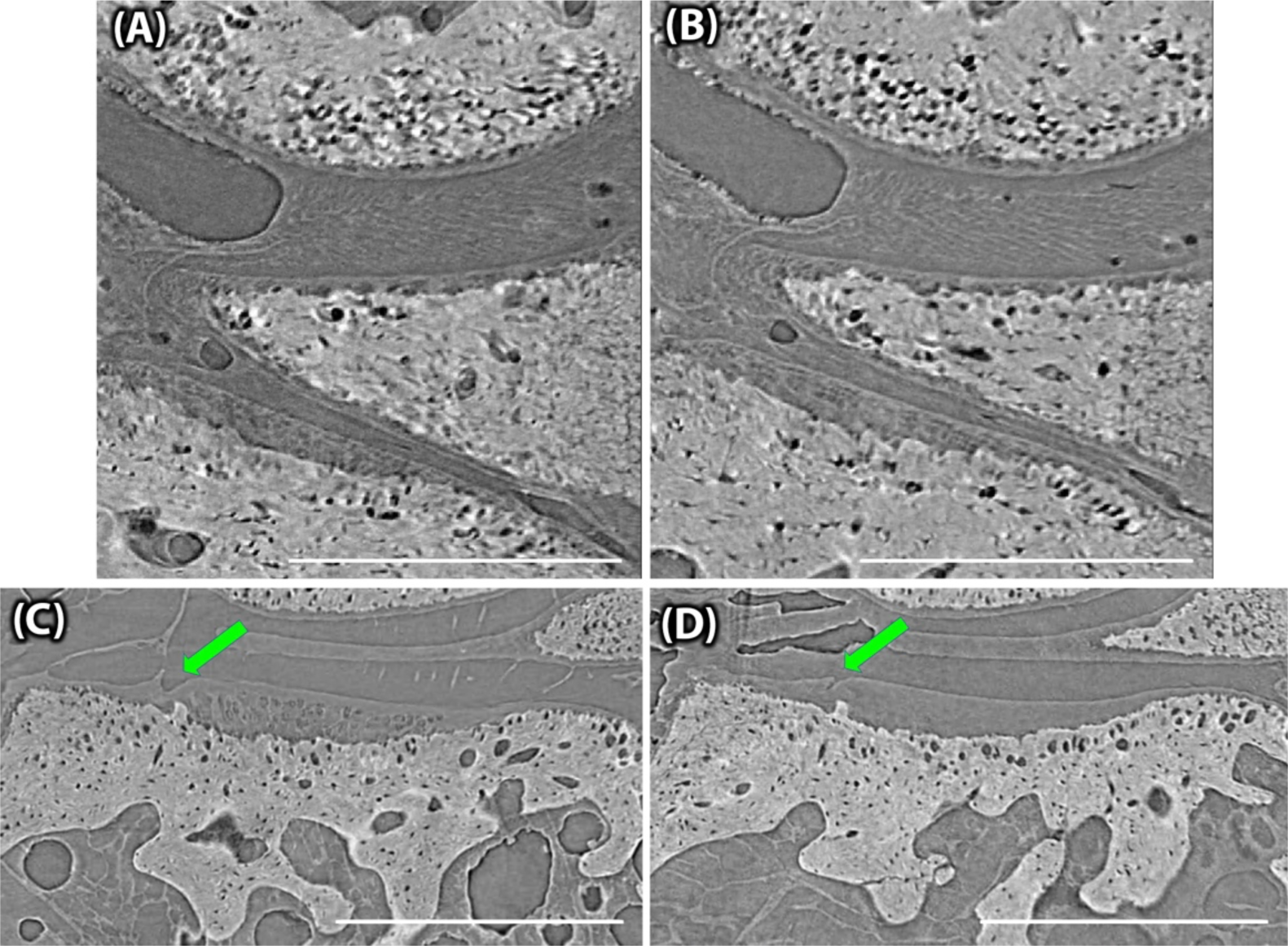
Propagation distance did not markedly affect phase contrast, but fixation impaired detail visualisation. Scale bars 500μm. (A) Fresh-frozen 20 week CBA mouse knee scanned at -20°C with propagation distances of 60mm and (B) 100mm. (C) 10 week male STR/Ort knee scanned with 100mm propagation distance, fresh frozen at - 20°C and (D) following room-temperature fixation in 4% PFA for 24 hours, then refrozen for scanning at -20°C. Note that tibial HAC cellular detail is visible in (C) but not (D), while a superficial HAC lesion (green arrows) also appears larger and more clearly visible in (C) than in (D).

### STR/Ort cartilage thickness is greater in most condyles, with early medial tibia differences

HAC was thicker in STR/Ort than age-matched CBAs in 16/24 compartments (all p<0.04). In contrast, HAC was thicker only in the posterior lateral femur of CBAs at 10wks (p<0.001, Table 1). At 20wks, however, the same compartment is the only region in which *no* significant strain difference in HAC thickness was observed.

Consistent with histological studies, HAC in the medial tibia of STR/Ort mice, in which OA first occurs, is already significantly thicker at 10wks (Fig. 3B)^(32)^. This was prominent posteriorly, where HAC thickness in STR/Ort mice is more than double that of age-matched CBA, and HAC thickness strain differences already exist across all condyles. This greater medial tibia HAC thickness in STR/Ort is maintained through to 36+wks.

**FIGURE 3:**
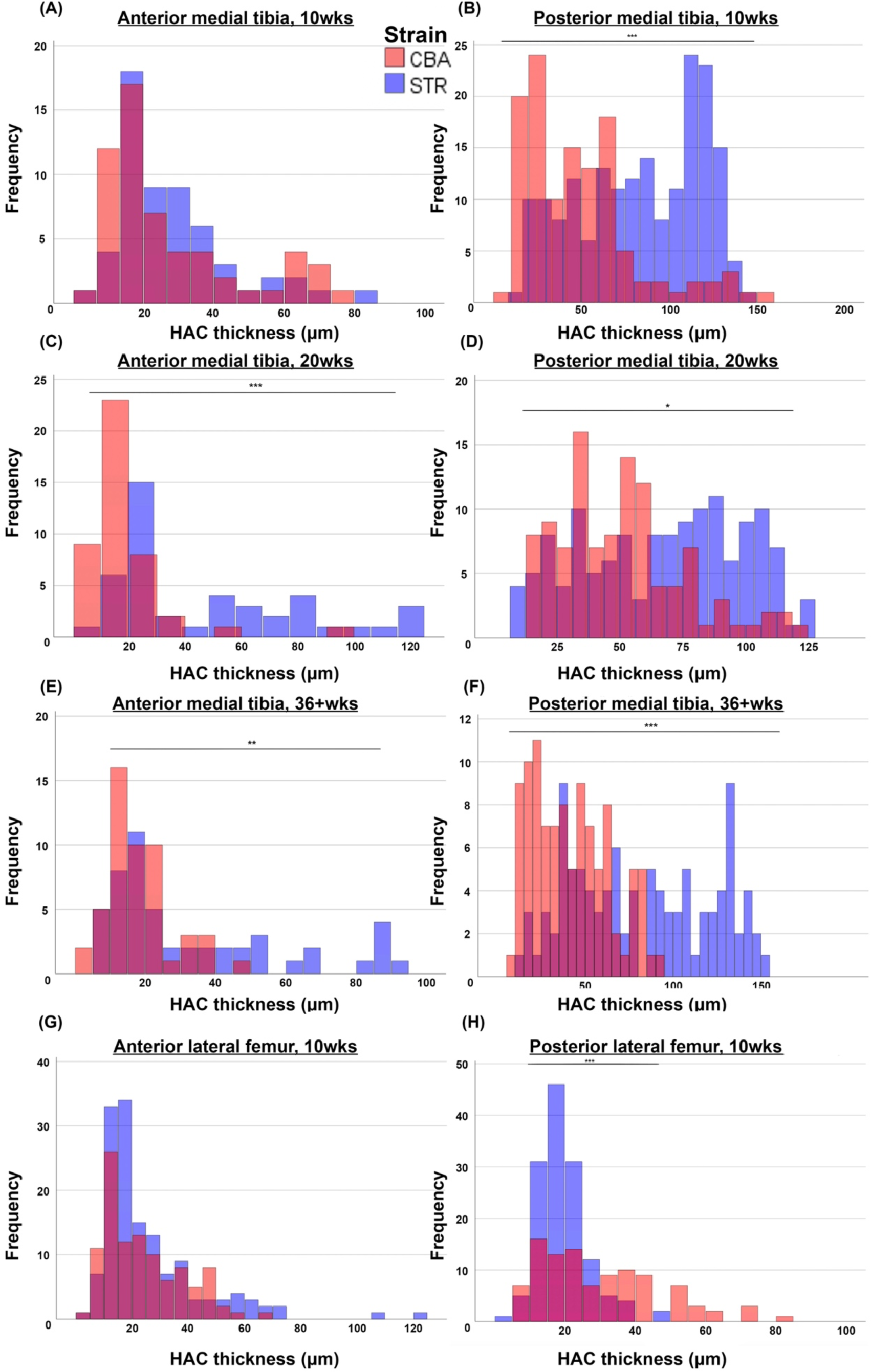
10 week old CBA mice only have significantly thicker HAC than STR/Ort mice in the posterior compartment of the lateral femur. (A) 10 week medial tibial anterior HAC thickness; (B) 10 week medial tibial posterior HAC; (C) 20 week medial tibial anterior HAC; (D) 20 week medial tibial posterior HAC; (E) 36+ week medial tibial anterior HAC; (F) 36+ week medial tibial posterior HAC; (G) 10 week lateral femoral anterior HAC; (H) 10 week lateral femoral posterior HAC. CBA mice (healthy controls) are represented in red, STR/Ort mice (osteoarthritic) in blue. Black significance notation indicates a significant difference between the two strains. * signifies p<0.05; ** signifies p<0.01; *** signifies p<0.001.

ACC was likewise significantly thicker in STR/Ort than age-matched CBA in 20/24 compartments (all p<0.03) at all ages. Intriguingly, strain differences in medial tibial HAC thickness at 10wks, restricted to the posterior condyle, were paralleled in ACC thickness (Table 2, Supplementary Fig. 3). Indeed, at no age or compartment were CBA found to have higher median ACC thickness than STR/Ort mice.

**TABLE 2:** Condylar compartments in which ACC thickness was measured, and the presence or absence of a significant difference between STR/Ort and CBA mouse strains. * indicates a significant difference between strains (p<0.05).

| Age (weeks) | Condyle | Anterior/Posterior | P-value for strain difference | Median CBA ACC thickness (µm) | Median STR/Ort ACC thickness (µm) |
| --- | --- | --- | --- | --- | --- |
| 10 | Lateral femur | Anterior | 0.016* | 60.0 | 71.4 |
|  |  | Posterior | < 0.001* | 60.3 | 83.7 |
|  | Lateral tibia | Anterior | 0.459 | 40.8 | 45.8 |
|  |  | Posterior | < 0.001* | 51.7 | 68.0 |
|  | Medial femur | Anterior | < 0.001* | 69.4 | 117.6 |
|  |  | Posterior | < 0.001* | 61.5 | 104.7 |
|  | Medial tibia | Anterior | 0.228 | 43.1 | 45.2 |
|  |  | Posterior | < 0.001* | 51.3 | 63.8 |
| 20 | Lateral femur | Anterior | < 0.001* | 69.1 | 90.5 |
|  |  | Posterior | < 0.001* | 59.7 | 86.2 |
|  | Lateral tibia | Anterior | 0.002* | 27.2 | 48.5 |
|  |  | Posterior | 0.023* | 56.8 | 60.6 |
|  | Medial femur | Anterior | < 0.001* | 66.9 | 112.1 |
|  |  | Posterior | < 0.001* | 63.7 | 99.5 |
|  | Medial tibia | Anterior | 0.002* | 44.9 | 56.2 |
|  |  | Posterior | < 0.001* | 54.6 | 69.6 |
| 36+ | Lateral femur | Anterior | < 0.001* | 65.2 | 92.6 |
|  |  | Posterior | < 0.001* | 56.0 | 79.9 |
|  | Lateral tibia | Anterior | 0.551 | 39.9 | 46.7 |
|  |  | Posterior | < 0.001* | 46.3 | 60.6 |
|  | Medial femur | Anterior | < 0.001* | 74.2 | 108.8 |
|  |  | Posterior | < 0.001* | 51.8 | 108.2 |
|  | Medial tibia | Anterior | 0.025* | 48.1 | 52.0 |
|  |  | Posterior | 0.178 | 60.1 | 63.7 |

As expected, total articular cartilage was thicker in STR/Ort than age-matched CBA mice in 20/24 compartments (all p<0.005). Again, strain thickness differences within the medial tibia were localised posteriorly at 10wks. As with HAC and ACC, significant whole-condyle differences in thickness exist by 20wks and persist at 36+wks (Table 3, Supplementary Fig. 4). Older STR/Ort mice retained a significantly thicker cartilage, at least in the non-lesioned regions, at 36+wks, and despite OA lesion development.

**TABLE 3:**
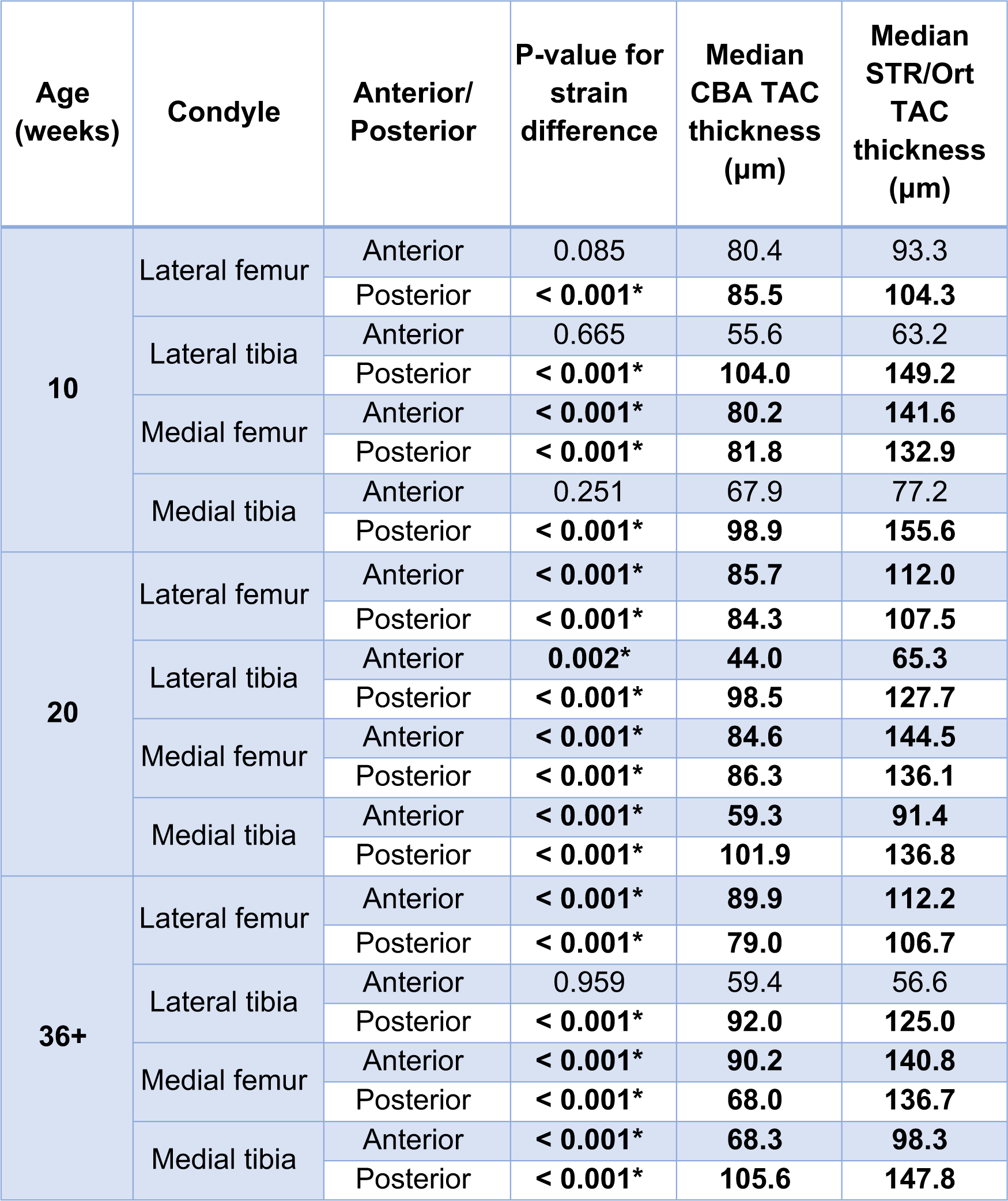
Condylar compartments in which total articular cartilage (TAC) thickness was measured, and the presence or absence of a significant difference between STR/Ort and CBA mouse strains. * indicates a significant difference between strains (p<0.05).

#### Early-life posterior strain differences in relative medial tibial HAC/ACC thickness

While our data verify both greater HAC and ACC thickness, it is possible their relative proportions are modified. Evaluation of *relative* HAC:ACC thickness (% total articular cartilage) detected significant strain differences in only 11/24 compartments (Tables 1-3). No significant strain difference was observed in anterior medial tibiofemoral joint regions at 10wks (p=0.083, Supplementary Fig. 5A) that was quantitatively most similar in the two strains, and unique as the *only* region without any significant strain differences in either absolute or relative HAC, ACC or total thickness (Tables 1-4).

**TABLE 4:**
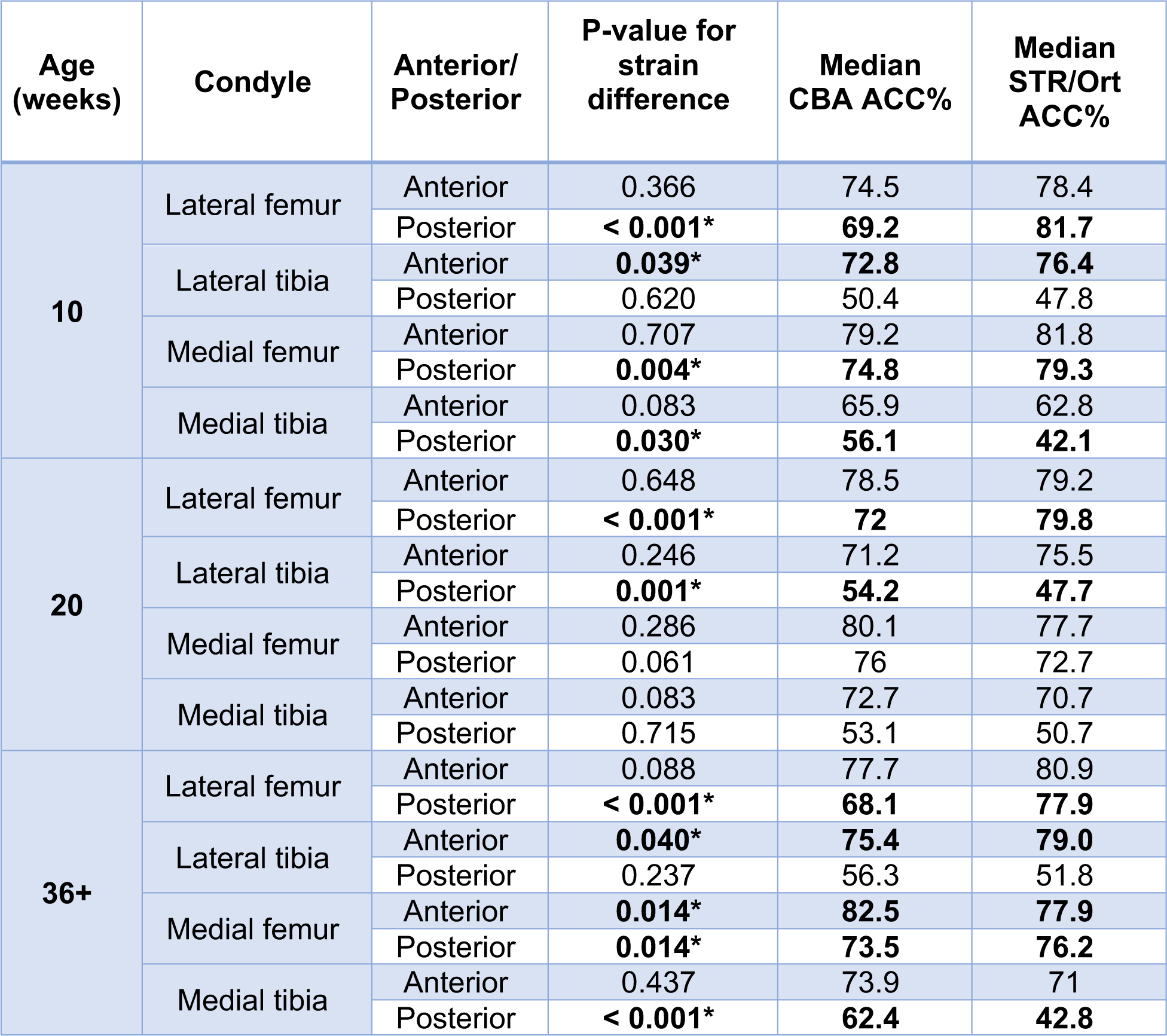
Condylar compartments in which percentage HAC and ACC thicknesses were measured, and the presence or absence of a significant difference between STR/Ort and CBA mouse strains. P-values shown are identical for both HAC% and ACC% strain differences. * indicates a significant difference between strains (p<0.05). Note that although only ACC% values are shown in the rightmost two columns, HAC% can be easily calculated as 100%-(ACC% value).

#### Both strains have thicker medial tibial posterior HAC/ACC at 10wks

HAC, ACC, total cartilage and relative ACC thickness were compared across anterior/posterior medial tibia. STR/Ort and CBA showed significant medial tibial antero-posterior asymmetry at 10wks, although both the *strength* of statistical significance, and *size* of biological difference between antero-posterior thickness values, were greater in STR/Ort than CBA (Table 5). The posterior medial tibia had thicker HAC and ACC than the anterior compartment in both strains at this age.

**TABLE 5:** Antero-posterior asymmetries exist in cartilage layer thicknesses within the medial tibial condyle of 10 week old STR/Ort and CBA mice. * = significant (p<0.05).

| | P-value difference anterior vs. posterior HAC thickness ($\mu\text{m}$) | P-value difference anterior vs. posterior ACC thickness ($\mu\text{m}$) | P-value difference anterior vs. posterior TAC thickness ($\mu\text{m}$) | P-value difference anterior vs. posterior ACC % | Median HAC thickness ($\mu\text{m}$) | | Median ACC thickness ($\mu\text{m}$) | |
| --- | --- | --- | --- | --- | --- | --- | --- | --- |
| **STR/Ort** | < 0.001* | < 0.001* | < 0.001* | < 0.001* | Anterior: | 25 | Anterior: | 45 |
|  |  |  |  |  | Posterior: | 87 | Posterior: | 64 |
| **CBA** | < 0.001* | 0.010* | < 0.001* | 0.013* | Anterior: | 19 | Anterior: | 43 |
|  |  |  |  |  | Posterior: | 43 | Posterior: | 51 |

#### Trans-zonal and ACC chondrocytes are larger in STR/Ort mice

Strain also affected chondrocyte lacunar volume (p=0.036), with greater hypertrophy in STR/Ort (1,351μm^3^) than CBA mice (946μm^3^). Post-hoc testing failed to confirm this strain difference in HAC (p=0.346, 1,383μm^3^ vs 1,217μm^3^ in STR/Ort and CBA), but it was pronounced for lacunae both in ACC (p=0.007, 1,737μm^3^ vs 947μm^3^, Fig. 5A) and trans-zonal regions (p=0.049, 933μm^3^ vs 675μm^3^, Fig. 5B). Similarly, lacunar surface area was significantly greater for STR/Ort than CBA (p=0.047).

#### Trans-zonal and ACC medial and lateral chondrocytes differ

An unexpected statistical interaction was seen between lacunar depth and condyle, with respect to chondrocyte volume. While volume of HAC and ACC lacunae did not differ significantly in the medial and lateral condyles (p=0.743 and 0.072), volume of trans-zonal chondrocyte lacunae was larger in medial than lateral condyles (p=0.021, 987 vs 769μm^3^, Fig. 4C). Significant depth:condyle interaction was also seen for surface area, which did not differ across condyles in HAC (p=0.803), while trans-zonal chondrocytes had greater surface area in medial (628μm^2^) than lateral tibia (p=0.020, 524μm^2^). Conversely, ACC lacunar surface area was smaller in medial than lateral tibia (p=0.033, 717μm^2^ and 810μm^2^).

**FIGURE 4:**
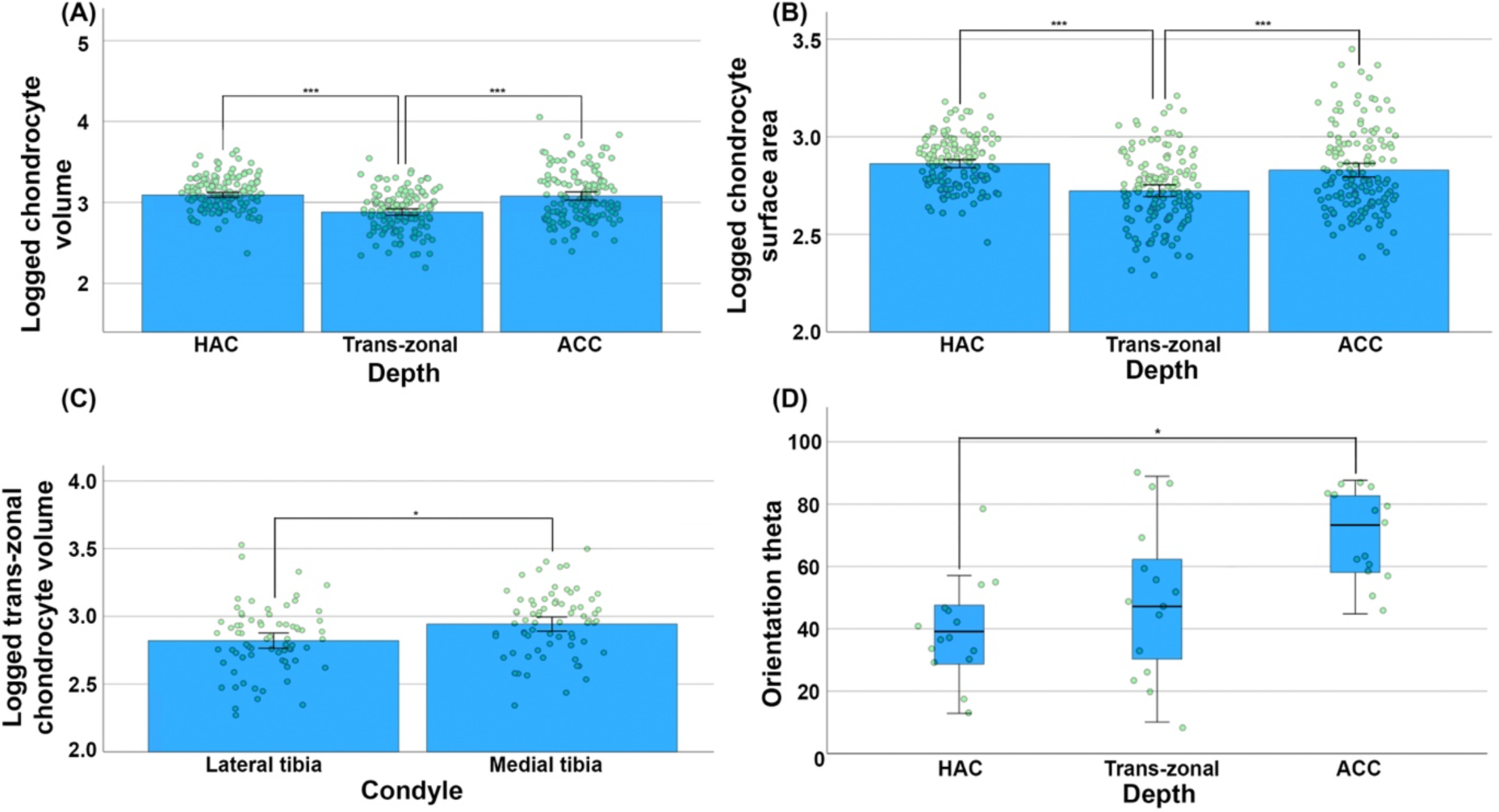
Chondrocyte size and orientation are region-dependent. (A) Trans-zonal chondrocytes are significantly smaller than either HAC or ACC chondrocytes. They also (B) possess lower surface area, specifically in the 10 week old STR/Ort lateral tibia. (C) Trans-zonal chondrocytes in the medial tibia are slightly larger on average than those in the lateral tibia. (D) HAC chondrocytes have lower orientation theta values, suggesting a relatively horizontal orientation, compared to ACC chondrocytes in 20 week old STR/Ort lateral tibias. * signifies p<0.05; ** signifies p<0.01; *** signifies p<0.001. Error bars show 95% confidence interval for the true mean.

### Chondrocyte orientation and sphericity are depth-influenced

In addition, 3D imaging facilitated pioneering evaluation of chondrocyte orientation (theta) which was significantly lesser in HAC than ACC in 20wk-old STR/Ort lateral tibias (p=0.031, 39.2° vs 73.3 °) suggesting that the longitudinal HAC chondrocyte axis is closer to a vertical orientation than a horizontal one, but vice versa for ACC chondrocytes. Chondrocyte sphericity was highly similar in HAC and trans-zonal lacunae (p=1.0, 0.770 vs 0.767). Both HAC and trans-zonal chondrocytes were, however, less spherical than those in ACC (both p<0.001, 0.812, Fig. 4D). This aligns with earlier findings that progenitor chondrocytes emerge in a flattened superficial HAC layer, becoming rounder as they traverse cartilage layers to hypertrophy in ACC.

#### Trans-zonal chondrocytes are smaller than ACC and HAC chondrocytes

Lacunar volume differed significantly in trans-zonal and deep ACC chondrocytes (p<0.001, 878 and 1,567μm^3^ respectively), and between trans-zonal and HAC lacunae (p<0.001, 878μm^3^ and 1,348μm^3^, Fig. 4A). Trans-zonal chondrocytes were smaller than those deeper or more superficial. Assessment of surface area confirmed trans-zonal chondrocytes are smaller than those by which they are surrounded, as their lacunar surface area (576μm^2^) was lower than in corresponding HAC and ACC (both p<0.001, 756μm^2^ vs 764μm^2^, Fig. 4B).

## Discussion

Our method enables unprecedented clarity in 3D HAC visualisation, and could be established as a gold-standard to measure ACC thickness and chondrocyte morphology (Figs. 1-2). Cryogenically contrast-enhanced sCT runs no risk of damaging cells through digestion isolation, staining or fixation artefacts, or wrinkled processing artefacts associated with wax-embedded sectioning. It allows thickness measurements in discrete increments without loss of slices. Our quantitative data reveal that HAC and ACC thickness are greater in STR/Ort than CBA mice in most regions throughout life, and is evident early on in the posterior medial tibia (Tables 1-2, Fig. 3). The capacity to measure 3D chondrocyte lacunar features across the intact endochondral interface has also shown trans-zonal chondrocytes to be larger in the medial than lateral joint compartment in both strains, but notably expanded in STR/Ort mice (Figs. 4-5). Intriguingly, our data also reveals shifting levels of chondrocyte orientation, sphericity and hypertrophy across the endochondral interface, with trans-zonal chondrocytes significantly smaller than chondrocytes in neighbouring layers.

**FIGURE 5:**
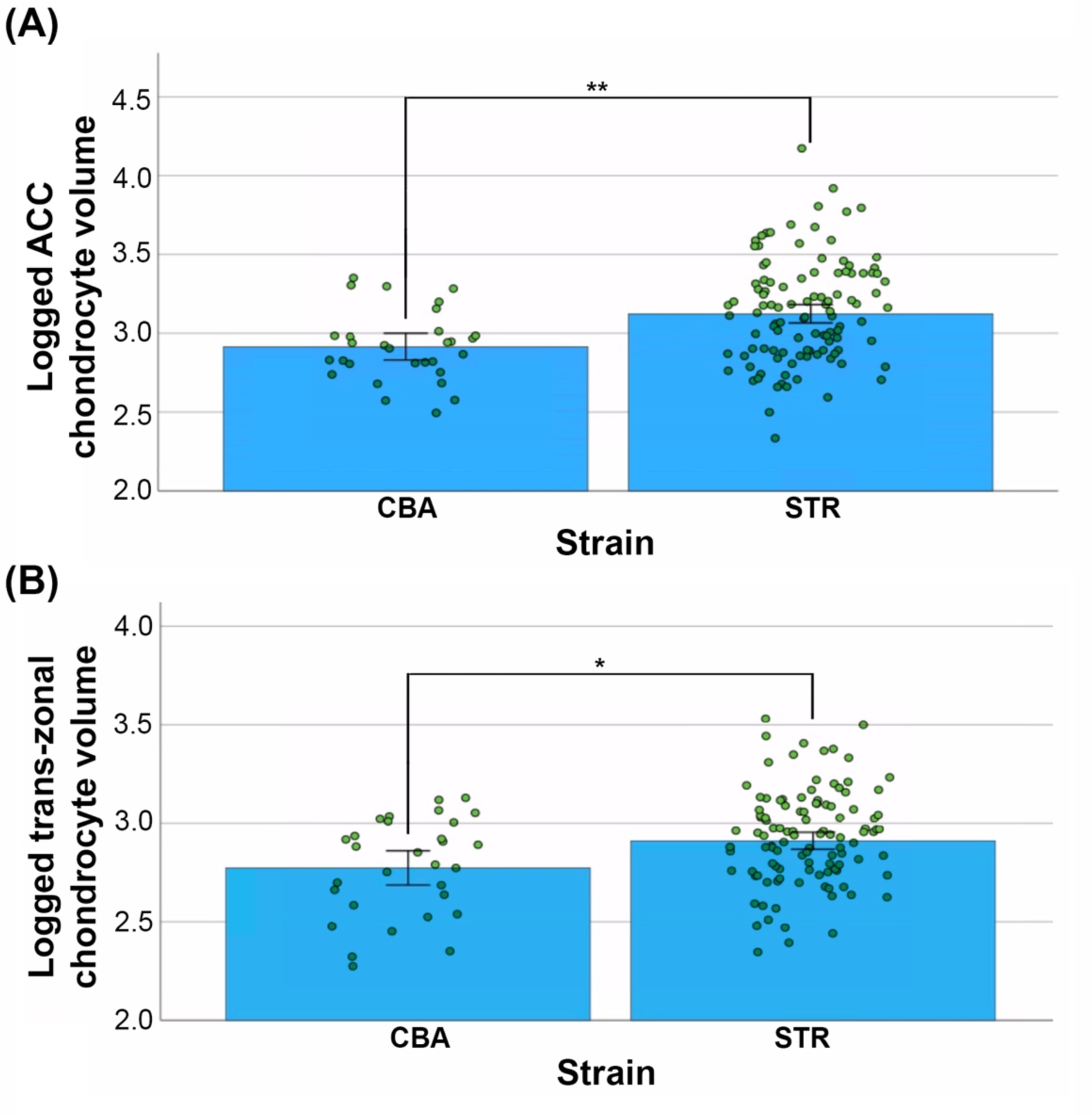
STR/Ort (A) ACC and (B) trans-zonal chondrocytes are larger than region-matched CBA chondrocytes. * signifies p<0.05; ** signifies p<0.01; *** signifies p<0.001. Error bars show 95% confidence interval for the true mean.

As this method makes HAC visible, with cellular detail (Figs. 1-2), 2D HAC thickness data could in future be spatially-linked to local 3D anatomical characteristics, such as medial tibial ACC; HAC volume; lesion location or growth plate thickness. This new visualisation method is superior to standard CT, revealing that: i) HAC is thicker across most condyles in STR/Ort (Table 1, Fig. 3H), ii) HAC and ACC are thicker in posterior than anterior medial tibial compartment at 10wks in both strains (Table 5), and iii) trans-zonal chondrocytes are a unique population with specific characteristics.

We found STR/Ort mice to have thicker HAC than CBA in most condyles, but young CBA HAC is thicker in the posterior lateral femur (Table 1). Thus, STR/Orts had thicker HAC, ACC and consequently, total cartilage than healthy CBA in most regions (Tables 1-3, Fig. 3). Significantly thicker HAC in posterior tibia of STR/Ort was maintained to late-stage OA (Table 1, Fig. 3F). This does not align with conventional narratives that age-related HAC thinning is an OA cause, or that JSW narrowing is likely a direct consequence of HAC lesions, and therefore a sensitive method to detect them. The exception is the posterior lateral femur, where HAC in young CBA is thicker (Table 1, Fig. 3H). Poulet et al. found that, with the knee in extreme flexion, the posterior lateral femur is the location where load-induced lesions are most readily generated in CBA in an *in vivo* joint loading model^(13, 44)^. Our findings therefore challenge current thinking by associating vulnerability to mechanical damage with regions of unusually thick, rather than ‘thinning’ cartilage. Indeed, our results align more with those of Omoumi et al.’s (2018) human research, which identified thicker HAC in the posterior medial femur of OA than non-OA patients^(17)^.

While STR/Orts and CBAs vary in cartilage composition in the posterior medial tibia, they were similar in the anterior compartment of this condyle at 10wks (Table 1, Fig. 3A). This was true for relative and absolute HAC and ACC (Table 4). This highlights a benefit of separate condylar lesion-scoring, as anatomical differences in the posterior condyle may reflect greater OA predisposition, and establish whether antero-posterior asymmetries in STR/Orts result in unfavourable load transfer onto a specific, vulnerable tibial region.

Mice have a crouched posture, with hindlimbs in near-constant flexion; body weight is likely to be transferred from femur to tibia primarily in the posterior knee. Given its load-bearing role, it is plausible mice have evolved thicker cartilage in the posterior tibial and femoral regions, consistent with our measurements (Table 5). Interestingly, Cohen et al. (2003) also found thicker cartilage in the posterior lateral femur of human knees, in OA and controls, compared to other femoropatellar regions^(45)^. More indicative of causality are thickness differences seen between STR/Ort and CBA; STR/Orts generally have thicker HAC and ACC, with a relatively thinner ACC layer as a proportion of total thickness. Spatially-mapping lesion location to determine whether they arise in regions with greater thickness prior to onset would enrich these findings and highlight possible cause/effect between overly-thick HAC and OA predisposition. It is plausible that thick HAC is due to pathological hydration changes early in OA^(46-48)^.

Human OA onset is commonly linked with thin cartilage, which may be due to thinning secondary to lesions, and therefore JSW measurements are decreased. The idea that articular cartilage thins with age, to predispose to OA, is challenged by Cohen et al. (2003) who find no significant difference in MRI-resolved cartilage thickness between 27-year-old volunteers and 56-year-old cadavers (both groups non-OA)^(45)^. It is conceivable these cadavers were exceptions to ‘usual’ cartilage thinning with age, or it had not yet onset. However, the burden of proof that thin, healthy cartilage is more OA-predisposed than thick, healthy cartilage, remains unsatisfied, and in STR/Orts, the opposite may be true.

Fewer differences were observed between strains in relative layer thicknesses. This implies evolutionary conservation of *relative* layer depth, supporting the hypothesis that ACC is equally important to cartilage integrity as HAC. This aligns with our finding that the posterior medial tibia compartment (OA onset site^(6, 49)^), was characterised in STR/Orts by less ACC at pre-OA (10wks). Finite element modelling, or indentation testing with digital volume correlation, may elucidate effects of this on weight transfer^(50)^.

Cryogenically contrast-enhanced sCT allowed for novel observations in chondrocytes spanning the tidemark, showing they are smaller than ACC and HAC chondrocytes (Fig. 4A). This accords with the notion that chondrocytes expand through cartilage to attain ACC hypertrophy. Prior to this finding, it was plausible chondrocytes achieve full hypertrophy while still partially in the deep zone, then being embedded in ‘size-appropriate’ lacunae. Our data imply that trans-zonal chondrocytes instead either become embedded in lacunae larger than the cells (inconsistent with our scans; Fig. 1I), or embed themselves in initially size-appropriate lacunae, later actively enlarged. This may explain local matrix metalloproteinase production associated with chondrocyte migration and ECM catabolism^(51, 52)^. As noted, quiescence of ACC chondrocytes does not necessarily equate to inactivity (Oegema et al. 1997)^(34)^.

Trans-zonal chondrocytes’ cell volume was also lower (Fig. 4A); surface area measurements were consistent with HAC and ACC chondrocytes having greater surface areas than trans-zonal counterparts. These data question dogma that chondrocytes expand through cartilage layers. Instead, our data indicate HAC chondrocytes undergo shrinkage (by ∼470μm^3^) *en route* to ACC (a new concept). Indeed, presumed inability to undergo voluntary shrinkage has contributed to hypotheses that only a specific chondrocyte subpopulation both expresses Col10a1 (the hypertrophic marker) whilst also remaining physically small, and only this cell undergoes secondary ossification centre osteoblastic transdifferentiation^(53, 54)^. Alternatively, it has been hypothesised that cell division is required for shrinkage in chondrocyte-to-osteoblast transdifferentiation^(53)^. Our data (Fig. 4) suggest instead chondrocyte shrinkage may be an innate characteristic.

Our data also indicate STR/Ort trans-zonal and ACC chondrocytes are larger than in CBA. Excessive hypertrophy in OA has been reported, and is linked with decreased capacity to form structurally-sound matrix and increased MMP-13 expression^(18, 33, 55)^. Larger STR/Ort lacunae described here may indicate exuberant hypertrophy, but may alternatively reflect greater fluid uptake, more cells in a stage of chondroptosis, differences in cytoskeletal organisation, or a combination^(18)^. The data showing trans-zonal chondrocytes were larger in medial than lateral tibial condyles is intriguing, given high STR/Ort OA predisposition medially. It is tempting to speculate that trans-zonal hypertrophy in the medial tibia contributes to OA, or reflects a pathological biomechanical loading environment causing hypertrophy, and hence OA. ACC chondrocytes being smaller in the medial than lateral condyle, supports the hypothesis of some condyle-specific difference in chondrocyte biology or environment, possibly linked to OA.

Finally, our imaging indicates that chondrocyte shape and orientation are depth-dependent. Significant chondrocyte orientation shifts at 20wks in STR/Ort mice is likely due to a larger sample size (Supplementary Table 2); conclusions about strain differences cannot be drawn. This shift in orientation is, however, informative of STR/Ort chondroskeletal ageing. Combined with findings relating to sphericity and volume, chondrocytes appear to become less vertically-oriented, more spherical, and (after brief trans-zonal shrinkage) eventually larger as they enter ACC.

## Conclusions

Cryogenically-enhanced phase contrast CT has clear benefits, revealing cellular detail in 3D as never before in murine HAC, which will allow further research regarding by-proxy measurements such as JSW and evaluation of soft tissue phenotypes, eg. Meniscal extrusion (uninvestigated in this model). This high-resolution imaging is therefore an exciting, novel method of examining samples while intact. With STR/Ort and CBA knees having been historically analysed by 2D histology, CT provides an opportunity to see whether 3D observations in these strains support, or even refute, conclusions from traditional methods. Association between excessively hypertrophic chondrocytes and OA has been reported, and our data affirm the presence of this link in STR/Ort mice^(6, 18, 56)^. Our data also surprisingly associate areas of OA predisposition with unusually thick, rather than thinner, cartilage, reminiscent of findings from Omoumi et al. (2018) in human patients^(17)^. In HAC, one interpretation is that some 10wk STR/Ort males are already in early OA, as hypothesised by Mason et al. (2001)^(57)^. Unusually-thick STR/Ort HAC may thus indicate cartilage over-hydration, reported before in human OA (Maroudas et al. 1985)^(46)^. Alternatively, thick HAC may be OA-vulnerable for another reason, eg. greater softness and correspondingly-increased exposure to pathological biomechanical forces; or increased hypoxia/anoxia due to greater distance from local vasculature^(58)^.

## Acknowledgements

Sincere thanks are due to Kaz Wanelik for his help with troubleshooting and image reconstructions, as well as to Nghia Vo for his help with reconstruction. Saurabh Shah was extremely helpful with the setup and use of the cold stage over multiple beamtimes, and thanks are due to the team who developed it, including Peter Rockett and Saurabh Shah. Phil Salmon and Mark Hopkinson have helped with interpretation of i3D data outputs, and Ruby Chang has given useful guidance on statistics.

## Authors

Lucinda Evans: Sample preparation, synchrotron scanning, image analysis, statistics, figures, original manuscript draft and further drafting.

Diana Vezeleva: Image analysis, statistics.

Andrew Bodey: Supervision, synchrotron experiments, image reconstruction, manuscript drafting.

Gowsihan Poologasundarampillai: Synchrotron scanning, image reconstruction, manuscript drafting.

Peter Lee: Supervision, cold stage provision, manuscript drafting. Andrew Pitsillides: Supervision, manuscript drafting.

