## supplementary for "Cryogenic non-invasive 3D X-ray phase-contrast imaging of unfixed, intact mouse joints reveals shifting chondrocyte hypertrophy across the endochondral interface"

**Supplementary Material: Cryogenic 3D X-ray phase-contrast imaging of unfixed, intact mouse joints reveals shifting chondrocyte behaviour at cartilage-endochondral interfaces**

Supplementary methods

Hand-segmentation of tibial hyaline articular cartilage (HAC), articular calcified cartilage (ACC) and trans-zonal chondrocytes at the tidemark

Hand-segmentation of individual HAC, ACC and trans-zonal chondrocytes was completed by a blinded investigator and fully reconstructed image stacks were opened in the coronal plane in CTAn (no increments). The central slice of each stack was identified (exactly midway between most anterior and most posterior slices) to show any tibial and/or femoral bone. To reduce analytical computing power required, stacks were then cropped to only 50 slices (81.25μm) on either side of this central region and the centre of the tibial condyle was identified (see Fig. 1B). The five chondrocytes closest to this centrepoint and fulfilling the following key criteria were identified in each of the three HAC, ACC and trans-zonal layers (n = 15 chondrocytes total per condyle, 30 chondrocytes per knee). Chondrocytes were fully hand-segmented on every slice (separation of 1.625μm), threshold-binarised, and subjected to 3D Individual Object (i3D) analysis in CT Analyser (Bruker, Belgium). Resulting data were checked to ensure that no values consistent with errors (eg. porosity or connectivity values other than 0) were contained.

### Figures and tables

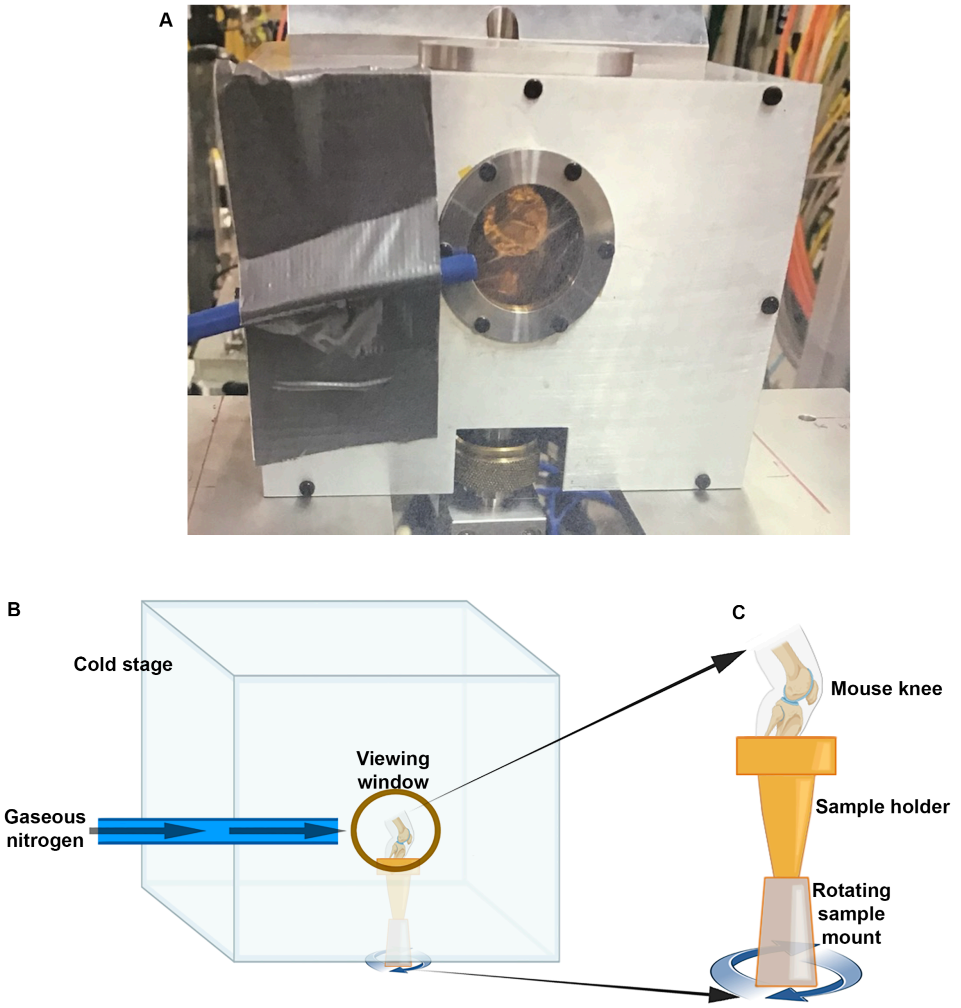


SUPPLEMENTARY FIGURE 1: *Frontal view of bespoke cold stage, showing its circular viewing window.* (A) Note also the blue tube taped in place; this fed a stream of room-temperature nitrogen gas over the Kapton tape in the window, illustrated in schematic shown in (B). (C) Enlarged diagram of sample mounting in cold stage. See also Guo et al (2017) for a more detailed sectional view of cold stage assembly[1]. (B-C) Created in BioRender and Adobe Photoshop 2023.

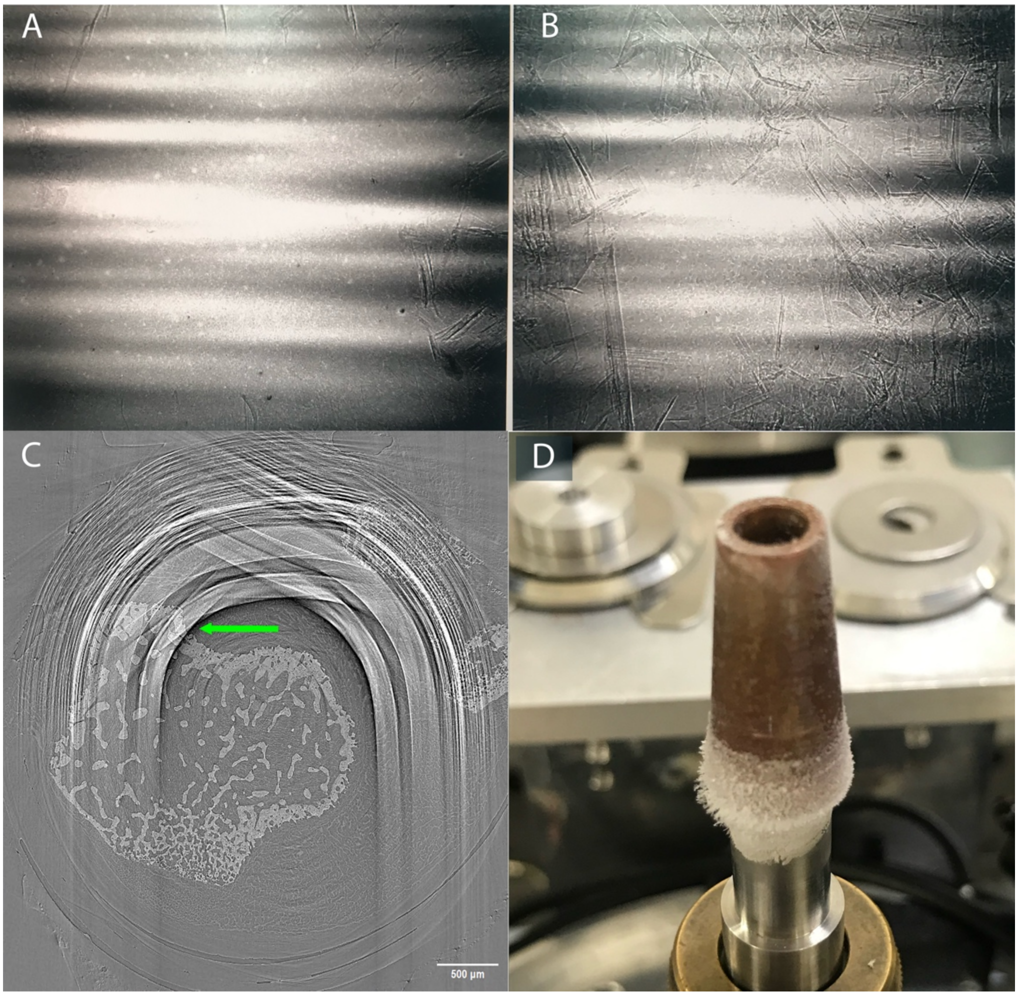


SUPPLEMENTARY FIGURE 2: *Cold-stage ice crystal formation, and its effect on scan outputs.* (A) Flatfield image (no sample) shortly after cold stage placement and cooling to -20$^{o}$C, prior to substantial ice crystal formation. (B) Flatfield view approximately 30 minutes later, during which time the stage was maintained at -20$^{o}$C, and numerous ice crystals had formed on the stage visualisation window. (C) The resulting extreme ring artefacts on an output image (9 week old male STR/Ort mouse tibia shown in transverse plane). The green arrow highlights a particularly problematic ring artefact, both sufficiently bright that it would incorrectly be labelled as ‘bone’ if threshold-based segmentation were used to isolate the femur from background, and sufficiently dark at the edge for true bone to be incorrectly labelled as background. (D) Ice crystals additionally formed outside the field of view around the brass sample mount, which needs to be able to rotate through 180$^{o}$ to obtain a scan. When the cold stage was placed on top of this holder, the ice could freeze it to the interior of the stage, preventing sample holder (and therefore, sample) rotation.

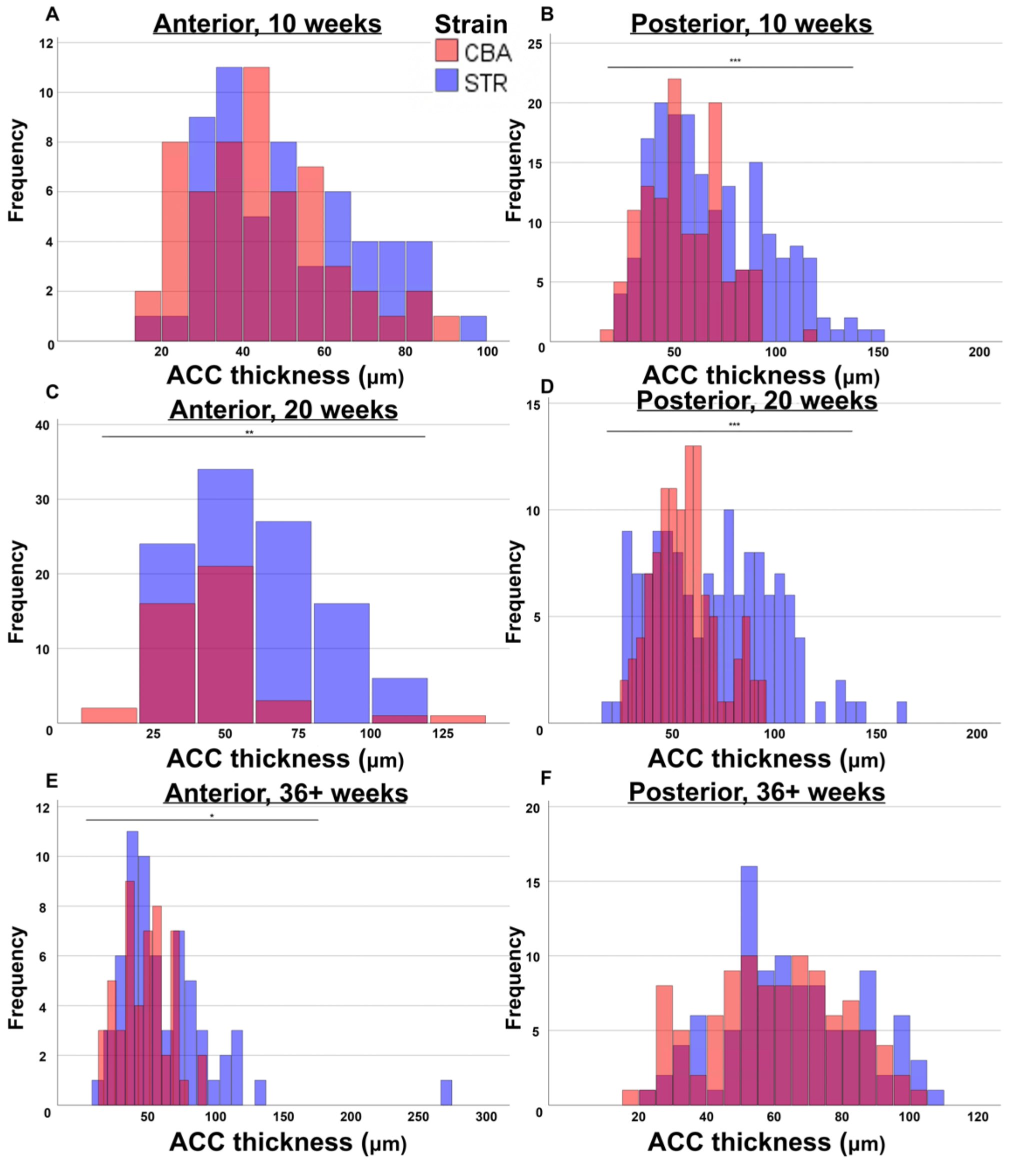


SUPPLEMENTARY FIGURE 3: *Medial tibial ACC thicknesses in STR/Ort and CBA mice at all age intervals.* (A-B), 10 week old mice; (C-D) 20 weeks old; (E-F) 36+ weeks old. (A), (C), (F) are anterior compartment; (B), (D), (F) are posterior compartment. CBA mice (healthy controls) are represented in red, STR/Ort mice (osteoarthritic) in blue. Black significance notation indicates a significant difference between the two strains. * signifies $p<0.05$; ** signifies $p<0.01$; *** signifies $p<0.001$.

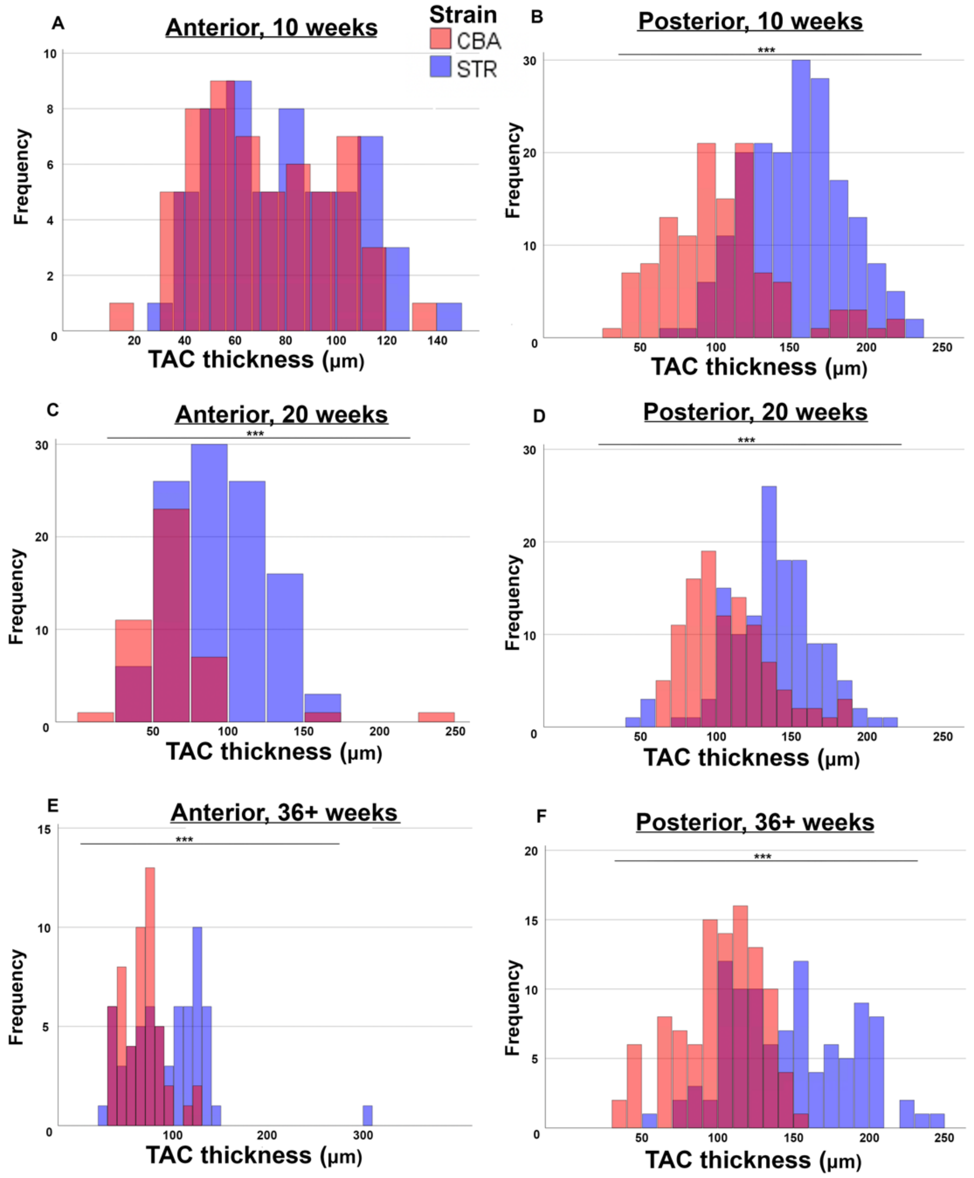


SUPPLEMENTARY FIGURE 4: *Strain differences in antero-posterior total articular cartilage (TAC) thickness exist in the medial tibial compartments of both strains throughout life, except in the anterior compartment at 10 weeks.* (A-B), 10 week old mice; (C-D) 20 weeks old; (E-F) 36+ weeks old. (A), (C), (F) are anterior compartment; (B), (D), (F) are posterior compartment. CBA mice (healthy controls) are represented in red, STR/Ort mice (osteoarthritic) in blue. Black significance notation indicates a significant difference between the two strains. * signifies $p<0.05$; ** signifies $p<0.01$; *** signifies $p<0.001$.

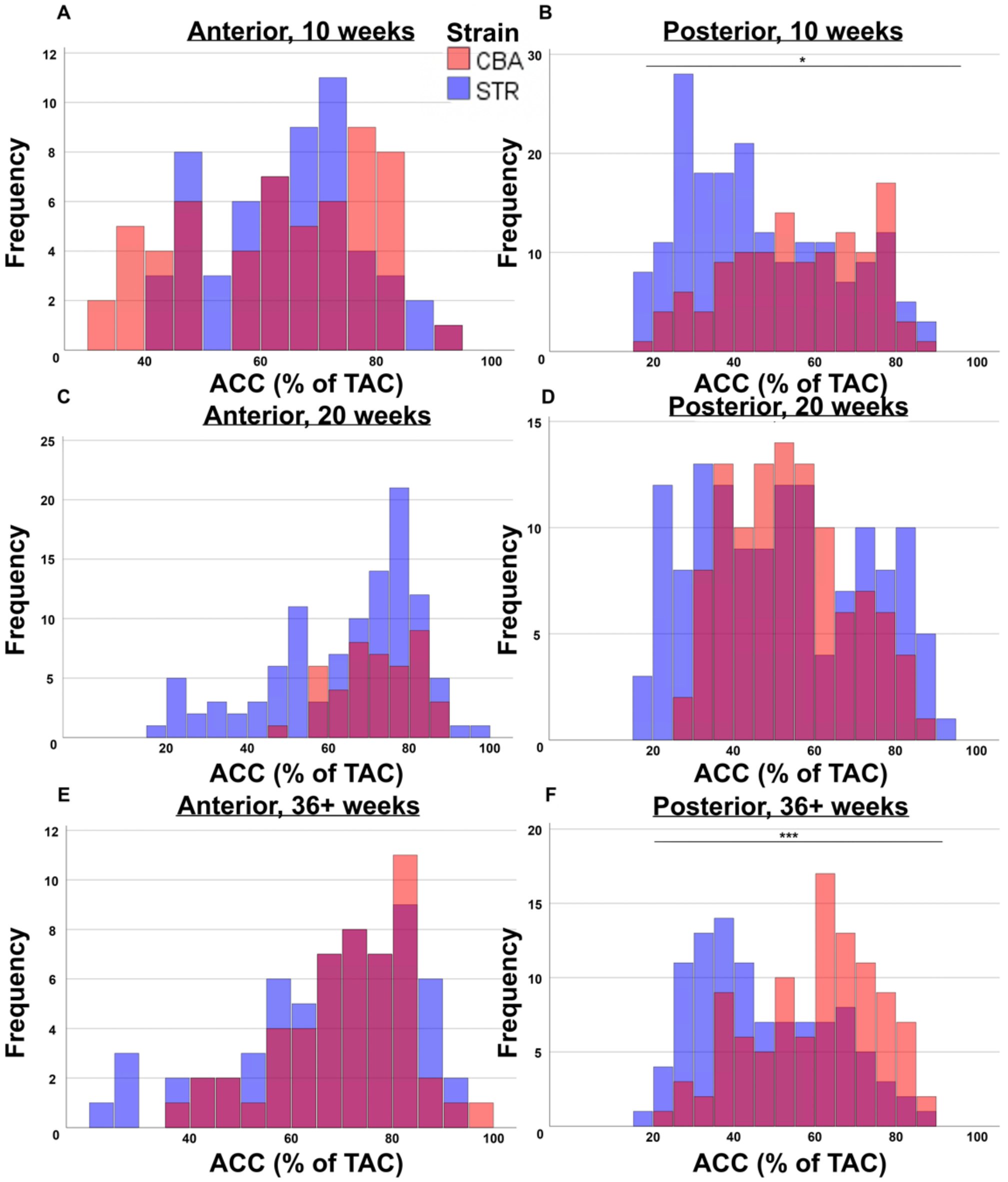


SUPPLEMENTARY FIGURE 5: *The percentage of medial tibial calcified cartilage differs between strains only in the posterior compartment – where CBAs have a relatively greater amount at 10 and 36+ weeks.* (A-B), 10 week old mice; (C-D) 20 weeks old; (E-F) 36+ weeks old. (A), (C), (F) are anterior compartment; (B), (D), (F) are posterior compartment. CBA mice (healthy controls) are represented in red, STR/Ort mice (osteoarthritic) in blue. Black significance notation indicates a significant difference between the two strains. * signifies $p<0.05$; ** signifies $p<0.01$; *** signifies $p<0.001$.

SUPPLEMENTARY TABLE 1: *Sample size of sCT-scanned mouse knee joints evaluated for cartilage thickness.*

| | **9-11 weeks** | **18-20 weeks** | **36+ weeks** |
|---|---|---|---|
| **STR/Ort** | 4 | 4 | 4 |
| **CBA** | 4 | 4 | 4 |

SUPPLEMENTARY TABLE 2: *Sample size of mouse tibias from which chondrocytes could be manually segmented for analysis.* Within each tibia shown in the table, 30 chondrocytes were segmented in total (5 each from the HAC, ACC and trans-zonal compartments, on both medial and lateral condyles: total n = 420). † = Note, the sample size of 2 tibias here is actually comprised from three different mice (one whole tibial plateau; one tibia in which HAC chondrocytes were visible only on the *medial* plateau; and one tibia in which HAC chondrocytes were visible only on the *lateral* plateau).

| | **10 weeks** | **20 weeks** | **36+ weeks** |
|---|---|---|---|
| **STR/Ort** | 6 | 3 | 2 |
| **CBA** | 2† | 0 | 1 |
